# Distinct N and C cross-feeding networks in a synthetic mouse gut consortium

**DOI:** 10.1101/2021.12.16.472894

**Authors:** Pau Perez Escriva, Tobias Fuhrer, Uwe Sauer

## Abstract

The complex interactions between gut microbiome and host or pathogen colonization resistance cannot solely be understood from community composition. Missing are causal relationships such as metabolic interactions among species to better understand what shapes the microbiome. Here, we focused on metabolic niches generated and occupied by the Oligo-Mouse-Microbiota consortium, a synthetic community composed of 12 members that is increasingly used as a model for the mouse gut microbiome. Combining mono-cultures and spent medium experiments with untargeted metabolomics uncovered broad metabolic diversity in the consortium, constituting a dense cross-feeding network with more than 100 pairwise interactions. Quantitative analysis of the cross-feeding network revealed distinct C and N food webs that highlight the two Bacteroidetes consortium members *B. caecimuris* and *M. intestinale* as primary suppliers of carbon, and a more diverse group as nitrogen providers. Cross-fed metabolites were mainly carboxylic acids, amino acids, and the so far not reported nucleobases. In particular the dicarboxylic acids malate and fumarate provided a strong physiological benefit to consumers, presumably as anaerobic electron acceptors. Isotopic tracer experiments validated the fate of a subset of cross-fed metabolites, in particular the conversion of the most abundant cross-fed compound succinate to butyrate. Thus, we show that this consortium is tailored to produce the anti-inflammatory metabolite butyrate. Overall, we provide evidence for metabolic niches generated and occupied by OMM members that lays a metabolic foundation to facilitate understanding of the more complex *in vivo* behavior of this consortium in the mouse gut.

**Importance:** This article maps out the cross-feeding network amongst 10 members of a synthetic consortium that is increasingly used as the model mouse gut microbiota. Combining metabolomics with *in vitro* cultivations, two dense networks of carbon and nitrogen exchange are described. The vast majority of the about 100 interactions are synergistic in nature, in several cases providing distinct physiological benefits to the recipient species. These networks lay the ground work towards understanding gut community dynamics and host-gut microbe interactions.

## Introduction

The mammalian gut microbiome is a complex community with thousands of bacterial species^1^ that affects many facets of host physiology, ranging from metabolism and development of the immune system to protection against pathogens ^2^. Extensive sequencing efforts categorized gut inhabitants and their genetic repertoire^3^, but fecal microbiome composition alone does not reveal the spatial and dynamic interactions between its members and with the host. These species interactions determine succession, stability, and resilience of a community^1^ and are the basis of causal relationships between microbiome composition and host physiology. Beyond correlative sequencing efforts, contemporary assessment of causal links is restricted to individual species^4^ or genes^5^. Understanding more complex behavior such as pathogen colonization, however, requires considering communities at large, which is hampered by technical limitations for *in vivo* studies. Recent *in vitro* studies demonstrated that elucidating the nature of pairwise interactions between community members can be used to predict assembly and dynamic behavior a community^6,7,8^. Such pairwise interactions can be neutral, negative for both (competition) or one partner (ammensalism), or positive for both (mutualism) or one partner (commensalism). The underlying basis may be physical^9,10^, quorum sensing^11^, toxins, competition for nutrients^12^, or metabolic cross-feeding.

To reduce the daunting complexity of natural systems, model communities of the gut microbiome with defined species composition have been used to colonize germ-free animals^13^. The primary focus of such models is to investigate complex phenotypes such as the interplay between the microbiome and host immune system or pathogen colonization resistance^14^. Recently, the Oligo-Mouse-Microbiota (OMM) consortium was introduced as a model for the mouse gut microbiome to study colonization resistance^15^. Composed of 12 natural murine isolates representing the five main gut phyla, it confers a higher colonization resistance towards the pathogen *Salmonella enterica* than the classical eight species altered Schaedler flora consortium^15^. Importantly, it is stable over time and reproducibly maintained in different animal facilities, rendering it an attractive model for the gut microbiome^16^. Although developed only recently, the OMM consortium has already helped to deepen our understanding of colonization resistance^15^, inflammation^17^, and development of the immune system^18^.

Generally, metabolic activities and interactions between species remain largely unexplored, even for these relatively simple, synthetic consortia. Analyzing extracellular metabolic changes upon growth in culture supernatants or in co-cultures revealed parts of a food web within the Altered Schaedler Flora consortium^19^. A first physiological characterization of microbial interactions within the OMM consortium reported primarily exploitative and interference competition during *in vitro* growth on culture supernatants^20^. From exometabolome changes in these cultures, the authors found substrate depletion profiles to correlate with growth inhibition, identified several species specific substrates and products, and singled out *Enterococcus faecalis* as the major determinant of community composition^20^, although is only a low abundant member of the healthy gut microbiome^21^. Actual cross-feeding of metabolites was hypothesized between *Clostridium innocum* and *E. faecalis*. Here we focus on unraveling cross-feeding systematically between all OMM species and ask whether such metabolic interactions could also be benefitial in nature rather than the reported competitive interactions, thereby contributing to community stability. Dynamic exometabolome changes during growth in complex medium and in culture supernatants of other consortium members revealed broad metabolic diversity amongst the OMM members that gave rise to a dense cross-feeding network with more than 100 pairwise interactions, where the most abundant *in vivo* members were the main providers. We unraveled two distinct food-webs of carbon and nitrogen sources that highlight Bacteroidetes as primary suppliers of C and Firmicutes as well as the Bacteroidetes *Muribaculum intestinale* as a provider of N containing compounds. The fate of several relevant cross-fed compounds was experimentally validated with isotopic tracing, allowing us to understand their metabolic fate within the community. We thus provide evidence for key metabolic niches that are generated and occupied by members of the OMM consortium and the individual roles of each member within it.

## Results

### Physiologic and metabolic diversity within the OMM consortium

To characterize physiology and secretion of metabolic products, each member of the OMM consortium was grown anaerobically in brain heart infusion broth supplemented with hemin, the vitamin K precursor menadione, and mucin as the key constituent of the gut mucus (mBHI)^22^. Representing less than five percent of the fecal bacterial load of mice carrying the OMM consortium^15^, the two minor constituents *Turicimonas muris* and *Acutalibacter muris* did not grow under these conditions. The other ten species achieved their maximum optical densities at 600 nm (OD) in mBHI within 25 hours (Fig. 1A). As the major constituents of the fecal community with up to 50%^21^, the Bacteroidetes phylum representatives *Bacteroides caecimuris* and *M. intestinale* exhibited similar lag phases of 4-8 hours and maximum ODs, but *M. intestinale* grew substantially slower (Table 1). The Firmicutes had shorter lag phases and displayed a broader range of maximum ODs and specific growth rates (Fig. 1A, Table 1). The mucus degrading *Akkermansia muciniphila* grew only to a low maximum OD and did not grow in the absence of mucin (Table S1), suggesting that mucin is its main carbon source, as shown previously^23^.

**Fig. 1.**
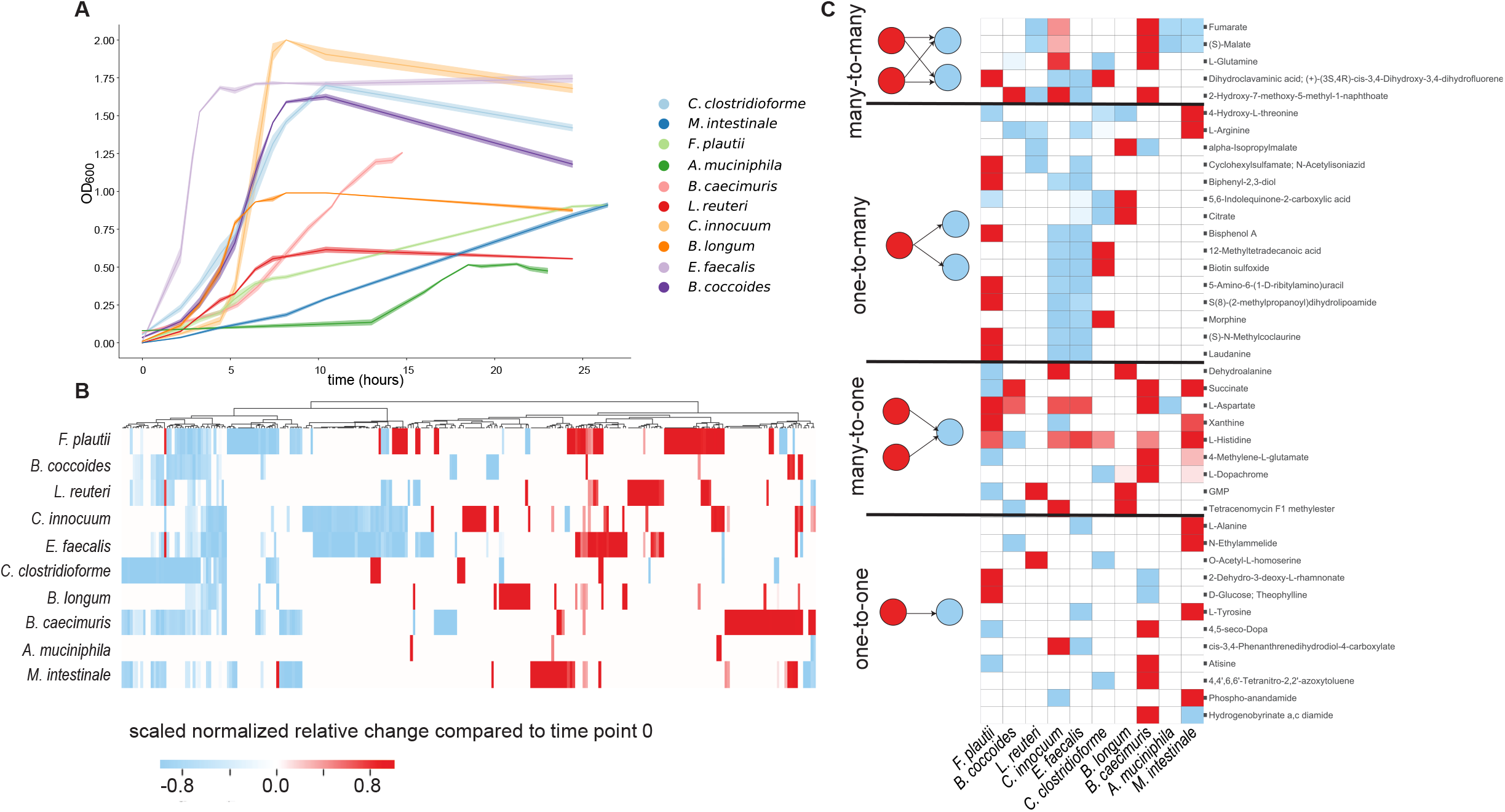
Exometabolome dynamics of OMM species in mBHI medium. **A.** Growth curves of 10 OMM species in mBHI medium. Shaded areas indicate the standard deviation from the mean (n=3-4 replicates). **B.** Metabolic footprint heatmap of all 10 OMM species during growth in mBHI medium. Secretion is indicated in red and consumption in blue. Intensities are scaled to +/− 1 by dividing each metabolite by the maximum observed change in abundance in all species. Hierarchical clustering was performed for metabolites, using Euclidian distances and centroid linkage. **C.** Heatmap describing the different types of potential interactions extrapolated from the consumption and secretion profiles in panel B.

**Table 1.** Physiological properties of the OMM members in mBHI. Cultures were grown in 10 ml of mBHI medium under anaerobic conditions in Hungate tubes.

| Phyla | Species | code | max OD | $\mu$ |
| --- | --- | --- | --- | --- |
| Firmicutes | *Flavonifractor plautii* | YL31 | 0.92 | $0.49 \pm 0.08$ |
| | *Blautia coccooides* | YL58 | 1.73 | $0.6 \pm 0.07$ |
| | *Lactobacillus reuteri* | l49 | 0.62 | $0.45 \pm 0.04$ |
| | *Clostridium innocuum* | l46 | 1.98 | $1.34 \pm 0.13$ |
| | *Enterococcus faecalis* | KB1 | 1.78 | $2.5 \pm 0.01$ |
|  | *Acutalibacter muris* | KB18 | - | - |
| | *Clostridium clostridioforme* | YL32 | 1.74 | $0.54 \pm 0.02$ |
| Actinobacteria | *Bifidobacterium longum* | YL2 | 1.01 | $0.91 \pm 0.04$ |
| Proteobacteria | *Turicimonas muris* | YL45 | - | - |
| Bacteroidetes | *Bacteroides caecimuris* | l48 | 1.18 | $0.47 \pm 0.02$ |
| | *Muribaculum intestinale* | YL27 | 0.93 | $0.39 \pm 0.03$ |
| Verrucomicrobia | *Akkermansia muciniphila* | YL44 | 0.32 | $0.09 \pm 0.01$ |

To characterize the metabolism of each species, we determined consumed and secreted metabolites by untargeted flow injection analysis time-of-flight mass spectrometry (FIA-TOFMS)^24^. Specifically, three to four biological replicates were grown per species, and 8-10 aliquots of culture supernatants were sampled throughout the growth phase of each and measured with FIA-TOFMS. 713 detected ions could be annotated to metabolites based on accurate mass, assuming single deprotonation, 268 had changing time profiles in at least one of the 10 investigated species (Table S2). Additionally, amino acids and short chain fatty acids were quantified with a targeted liquid chromatography LC-MS method. The 29 metabolites consumed by at least half of the species were primarily organic acids such as pyruvate and 2-oxobutanoate, amino acid derivatives such as 4-phospho-L-aspartate, N-succinyl-L-citrulline and 5-hydroxy-L-tryptophan, and a few micronutrients such as ascorbate (Table S2, Fig. S1A, S1C).

Despite their abundance in mBHI, surprisingly none of the amino acids was consumed by the majority of the members (Fig. S2). Besides serving as precursors for biomass, amino acids can also be used as an energy source, for example in Stickland fermentations, where one amino acid serves as an electron donor and another as an acceptor by forming carboxylic acids^25^. Many gut microbes are capable of Stickland reactions that may play a role in cross-feeding in the gut^26^. Several OMM members have the capacity to degrade arginine, alanine, glycine, leucine, or aspartate by Stickland fermentation^20^. The observed degradation of asparate by *A. muciniphila* and alanine by *E. faecalisi* might thus be explained by Stickland fermentation (Fig. S2). Although all OMM members could potentially degrade arginine via Stickland reduction, only four members consumed it under our conditions (Fig. S2). Other types of amino acid degradation were, for example, seen for lysine and histidine that were consumed by *Flavonifractor plautii* and *Blautia coccoides*, respectively, the only OMM members able to degrade these amino acids (Fig. S2, Table S2)^27^. Lysine degradation might be relevant *in vivo* for this consortium since its degradation by *F. plautii* yields two of the three classical short chain fatty acids, acetate and butyrate. Although nine species encode L-serine dehydratase orthologs that can catalyze the degradation of serine to pyruvate, only the fast-growing *E. faecalis* consumed serine in larger amounts and to a smaller extent *C. clostridiforme* and *F. plautii* (Fig. S2, Table S2).

Typical end products of fermentation such as short chain fatty acids^28^ and amino acid derivatives^29^ were produced by some species. In particular acetate was secreted by most consortium members (Fig. S2). The short chain fatty acid butyrate that is used by enterocytes as an energy source^30^ was produced by *C. clostridiforme, F. plautii* and *C. innocuum* (Fig. S2), where the latter two have the genetic repertoire for its production from sugars^27^. Propionate was secreted by *A. muciniphila* and the two Bacteroidetes members with succinate pathway genes, the only propionate production pathway from carbohydrates known for this phylum^31^. Accumulation of amino acids, most likely from peptide digestion^32^, was seen for the two Bacteroidetes members and several Clostridia (Fig. S2). Amino acid fermentation products such as isopropyl-malate, 4-aminobutanoate and 4-methyl-oxopentanoate and 3-methyl-2-oxobutanoic acid were secreted by *C. innocuum, B. longum* and *M. intestinale* (Table S2). While many metabolites were secreted by several species, not a single one was secreted by all (Fig. 1B). The large number of metabolites (90, 33% of all changing metabolites) secreted by only one species and several taxon-specific metabolites suggests a broad metabolic diversity (Fig. S1B, S1D). With 52 secreted metabolites (19% of all the metabolites detected to change over time), *F. plautii* was not only the main producer in the consortium but also the unique source of 23 metabolites (Fig. S1D).

To start mapping out the metabolic interaction network from the consumption and secretion patterns in monocultures, we selected for metabolites that were secreted by at least one member of the consortium and consumed by one or more members. From the 268 compounds with dynamic profiles, we predict 15 as one-to-many, 12 as one-to-one, 9 as many-to-one, and 5 as many-to-many cross-feeding interactions (Fig. 1C). In total, 41 metabolites were secreted by at least one member and consumed by at least one other, hence are potentially cross-fed.

### The OMM metabolic food web is highly connected

To obtain more direct evidence for cross-feeding and to capture interactions through secreted metabolites that were not already present in mBHI, we performed systematic pairwise cultivation experiments. For this purpose, cell-free culture supernatants of all ten species were harvested at maximum OD in mBHI medium. These supernatants (i.e. spent media) were mixed at a ratio of one to one with fresh mBHI, to ensure some bacterial growth, and inoculated with each of the other nine species in duplicates. To assess spent media influence on consumers, we compared the maximum OD obtained in the spent medium to the one obtained in undiluted mBHI (Fig 2A). In one case the maximum OD of the consumer was even 10% higher than in undiluted mBHI; *i.e*. when *B. caecimuris* was grown in *A. muciniphila* spent medium (Fig. 2A). More generally, six out of the eight consortium members grew to nearly the same density as in pure mBHI on *A. muciniphila* spent medium, suggesting that *A. muciniphila* makes breakdown products of the complex glycoprotein mucin available to the community. Conversely, *A. muciniphila* grew poorly in most spent media except those of *L. reuteri* and *B. coccoides*,presumably because these two species consumed less of *A. muciniphila’s* main nutrient source mucin. *L. reuteri* does not have the genetic repertoire for mucin degradation while *B. coccoides* does (Table S4). Hence, the latter may still degrade mucin in mBHI but provide other metabolites to *A. muciniphila*. While the attained maximum ODs were generally lower than in undiluted mBHI, most of them were higher than the half maximum OD that one would expect from a one to one dilution (Fig. 2A), indicating consumption of additional nutrients from culture supernatants or non-overlapping nutrient preferences between the two species. Besides the above nutritional benefit of *A. muciniphila* culture supernatant, a particularly beneficial combination was seen between *F. plautii* and *M. intestinale*. Five cultures reached lower maximum ODs than expected from a one to one dilution and seven cultures did not grow at all, mostly *M. intestinale*, suggesting either competition for essential nutrients or secretion of inhibitory metabolites. Investigating the OMM species in pure spent media, a parallel study found mainly growth inhibition^20^, possibly as a consequence of nutritional competition. The high frequency of positive interactions in our experiments was probably caused by mixing spent medium one to one with undiluted mBHI, which avoids growth inhibition through exhaustion of essential metabolites.

**Fig. 2.**
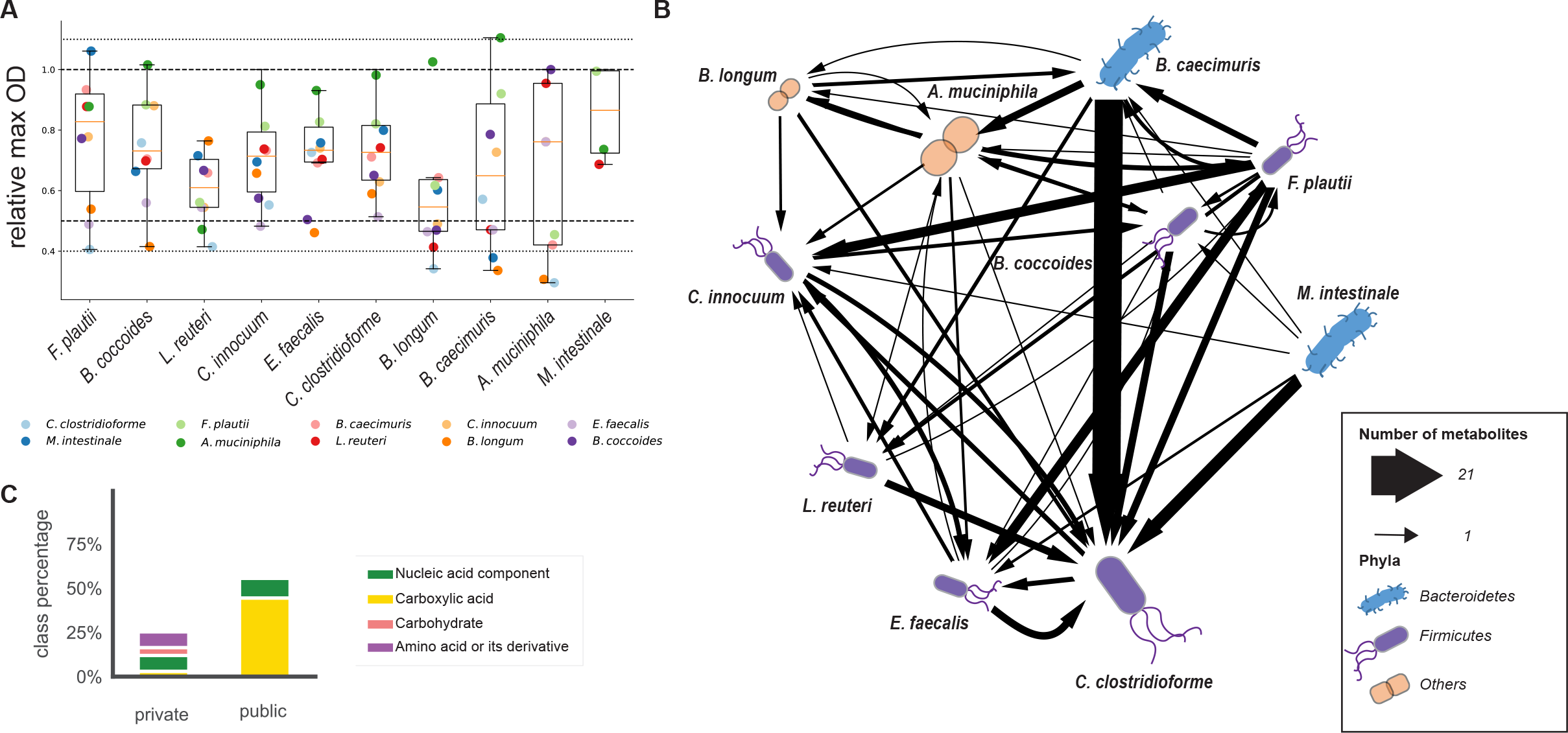
Metabolite cross-feeding among OMM members in mBHI spent medium. **A.** Maximum OD of OMM species during growth in a spent medium mixture of 50% culture supernatant and 50% mBHI. Data points are the mean of duplicate measurements (Table S2). The species-origin of culture supernatant is indicated by the color of the points. The relative maximum OD achieved by each of the other species is given on the y-axis, relative to the maximum OD achieved in fresh mBHI. Relative maximum ODs of 1.1, 1, 0.5 and 0.4 are indicated by dotted lines. **B.** Metabolite interaction network of OMM species inferred from spent medium experiments. Metabolites secreted by a producer in fresh mBHI and consumed in spent media by a second species were classified as cross-fed. Consumed metabolites in spent medium experiments were identified by filtering all decreasing annotated ions, either based on a significant correlation with the culture OD over time (Pearson correlation coefficient<-0.7, p-value<0.05) or a significant goodness of linear or exponential fit (R2>0.7, p-value<0.05). **C.** Relative metabolite class distribution of public and private cross-fed compounds. Percentages were calculated from the total numbers of metabolites within a class divided by the total number of metabolites, including the ones without a specific class associated.

Cross-fed metabolites were identified from dynamic pattern of extracellular metabolites during growth in the spent media experiments with FIA-TOF MS. Similar to fresh mBHI, 216 annotated metabolites exhibited changing time profiles across all experiments. In total, 76 metabolites were secreted in fresh mBHI and consumed in spent media, 31 of which were already hypothesized to be cross-fed in the experiments with fresh mBHI (Fig. 1C, Table S2), providing evidence for 146 metabolic cross-feeding interactions (Fig. 2B, Table S5). For example, the organic acid succinate was cross-fed seven times as it was produced by several members (Table S5, Fig 1C). The largest number (21) of metabolic interactions was observed for *C. clostridioforme* when grown in the spent medium of *B. caecimuris*, although without apparent effect on the maximum OD (Fig. 2A). With up to 57 consumed metabolites, *C. clostridioforme* was the most promiscuous species and *M. intestinale* as the other extreme did not consume any of the detected metabolites that were secreted by other consortium members (Figure 2B). The small growth improvement of *F. plautii* in *M. intestinale’s* medium in the absence of detected cross-feeding suggests either the presence of a not detected metabolic interaction or the existence of an advantageous non-metabolic interaction. For a more systematic scoring of consumers and producers, we determined the ratio of consumption versus secretion interactions. Representing more than 50% of the OMM consortium in the mouse cecum and colon (Fig S3)^15^, the two Bacteroidetes *B. caecimuris* and *M. intestinale* had the lowest consumption-secretion ratio. The second most abundant *in vivo* Firmicutes - *F. plautii -* had the third lowest ratio (Table S6). Thus, *in vivo* abundant OMM members appear to have mainly a provider role within the cross-feeding network through a wide range of secreted compounds. The number of consumed metabolites was highly correlated with genome size (0.70), and even higher when considering only cross-fed metabolites (0.80), consistent with previous observations that specialist bacteria have smaller genomes than generalist bacteria^33,34^ (Table S6).

The 76 cross-fed metabolites include six carboxylic acids, six amino acids or derivatives thereof, seven nucleic acids, and three carbohydrates (Table S7). These metabolites may be either a private or public good; *i.e*. consumed by two or more members, respectively (Fig. 2C). As typical fermentation end products, carboxylic acids were significantly enriched among the public goods (Fisher-exact test, p-value = 0.001). For example, the organic acid succinate was cross-fed seven times. Of special interest are malate and fumarate, both secreted by *B. caecimuris* and *C. innocuum*, that can be used as electron acceptors for respiration in anaerobic environments^35^. These public goods were consumed by *L. reuteri, E. faecalis, A. muciniphila* and *C. clostridioforme*. The presence of several consumers for electron acceptors is potentially relevant for colonization resistance to *Salmonella* infections, which has been shown to require them during initial gut colonization^36^. As potential nitrogen or carbon sources^29^, amino acids were predominant among the private goods. For example, histidine was consumed only by *B. coccoides*, the genome of which encodes a degradation pathway that produces glutamate and formate from histidine. Thus, histidine could be used by *B. coccoides* as a biomass precursor, a nitrogen source, or to produce formate as an electron donor^37^. Another private good example is cysteine provision by several members to *C. clostridioforme* (Table S2). Overall, we thus provide evidence for a dense network of 146 cross-feeding interactions between the ten OMM members (Fig. 2B), mainly through carboxylic, amino acids and nucleobases by the three providers *B. caecimuris, M. intestinale*, and *F. plautii*.

To assess the relevance of the so far mapped interaction network and to unravel the underlying metabolism, we next quantified absolute metabolite concentrations in the above spent and fresh media experiments using a targeted LC-MS method covering 15 of the cross-fed compounds and additional compounds that were expected to be cross-fed from the behavior in fresh media (Fig. 1C). Concentrations of secreted compounds ranged from low micromolar to millimolar and were generally higher for public goods. As expected, fermentation end products accumulated to high concentrations with succinate being by far the most abundant cross-fed metabolite, reaching 14.7 ± 0.5 mM in *B. caecimuris* spent medium. As another public good, the nucleobases xanthine and hypoxanthine were secreted up to about 1 mM. Among the private goods, the metabolite cross-fed at the highest concentration was histidine (~0.56 mM)(Fig. 3).

**Fig. 3.**
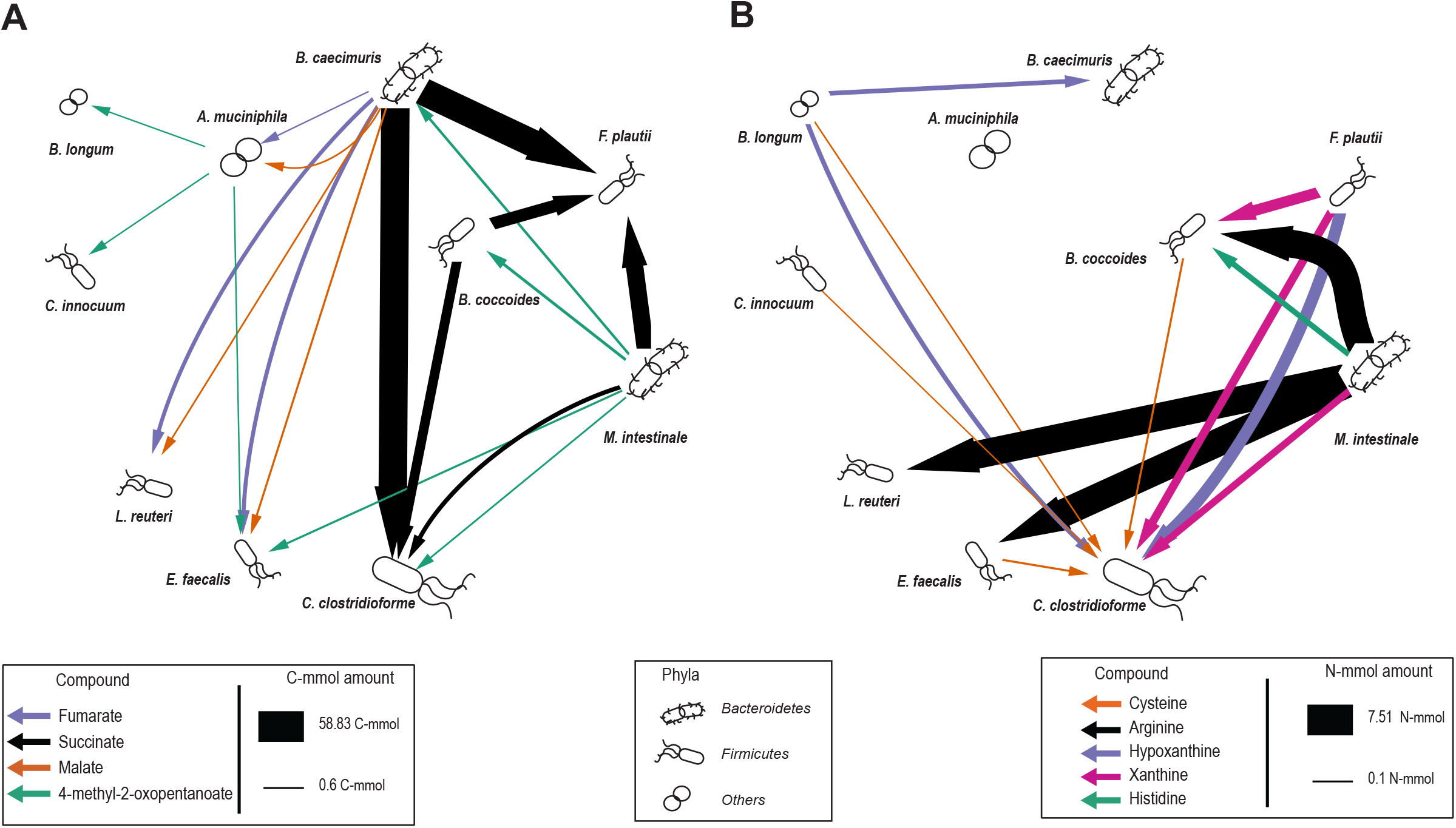
Carbon and nitrogen interaction networks of the OMM consortium. Interactions were inferred from growth experiments in a mix of 50% complex mBHI and 50% spent media of each OMM species. **A.** Compound specific OMM carbon interaction network. **B.** Compound specific OMM nitrogen interaction network. For both networks the amount of cross-fed compounds containing carbon or nitrogen was quantified and multiplied by the number of carbon or nitrogen atoms per molecule, respectively. Only compounds above 0.1 C-mmol or N-mmol exchange are displayed. Bacteria that represent more than 10% in the community are shown in larger size^21^.

When calculating the C and N-mol mass balance of cross-fed metabolites from the quantified consumption and production profiles, two rather distinct food webs emerge for carbon and nitrogen (Fig. 3). While Bacteroidete*s* were the main provider of C compounds, with succinate accounting for the major carbon flow (Fig. 3A), species from different phyla contributed to N flow, including *F. plautii, B. longum and M. intestinale*, the latter providing the major N flow through arginine (Fig. 3B). Of note, several amino acids increased over time during growth of *M. intestinale* (Fig. S2), most likely derived from peptide digestion. Somewhat surprisingly only arginine and histidine were found to be consumed by other species. Although many metabolites are cross-fed (Fig. 2B), the interaction between any two members was dominated by single C- and N-containing metabolites. The major mass flow of C was mediated by succinate and to a lesser extent by malate and fumarate (Fig. 3A), and for N by arginine followed by the purine degradation products hypoxanthine and xanthine, and histidine (Fig. 3B). Arginine catabolism was previously reported for lactic acid bacteria^38^ such as *L. reuteri* and *E. faecalis* via the 3 step arginine deaminase system for energy generation with ornithine as a side product^38^. Both *L. reuteri* and *E. faecalis* contain all necessary genes, including the arginine-ornithine antiporter ArcD that couples arginine uptake to ornithine secretion (Table S8, S9). Consistently, we observed ornithine secretion in both *L. reuteri* and *E. faecalis* in mBHI (Table S10) and that this secretion was greater when grown in the spent medium of the arginine-producing *M. intestinale*. This cross-feeding interaction might also be relevant for the host because ornithine production by *Lactobacillus* has been shown to contribute to the maintenance of a healthy gut mucosa^39^.

Overall, succinate, malate and fumarate dominated cross-feeding in the C-network and the amino acids arginine and histidine as well as the nucleobases xanthine and hypoxanthine the cross-feeding in the N-network. In molar terms, *B. caecimuris* and *M. intestinale* were the predominant providers of C and N, and *F. plautii* the predominant C consumer and at the same time a relevant N provider. While microbial cross-feeding of succinate has been previously reported in the gut^40^, the extent to which malate and fumarate or xanthine and hypoxanthine are cross-fed has not been characterized yet.

### Supplementation reveals physiological benefits of cross-feeding

With nine compounds being cross-fed at more than 100 μM (Table S11), we next investigated the physiological relevance of this C and N flux between species. Cultures of consuming species were grown in mBHI medium separately supplemented with 10 mM each of the 9 compounds to determine specific growth rate and maximum OD. While most supplemented metabolites did not affect either of these physiological parameters, as might be expected in a rich complex medium like mBHI, malate almost doubled the growth rate of *L. reuteri* and *A. muciniphila* and fumarate had an even more dramatic effect on *L. reuteri* (Fig. 4A). Supplementation with N-containing compounds only affected the maximum OD; i.e. a 10-20% improvement of *B. coccoides* and *C. clostridioforme* by xanthine and the former also by histidine. Although arginine has been reported as a C and N source^41^, supplementation did not improve growth of *L. reuteri* or *E. faecalis*. Supplementation with the amino acid cysteine had a drastic negative impact on *C. clostridioforme* (Fig. 4A), although it was consumed during growth in spent media (Fig. 3B).

**Fig. 4.**
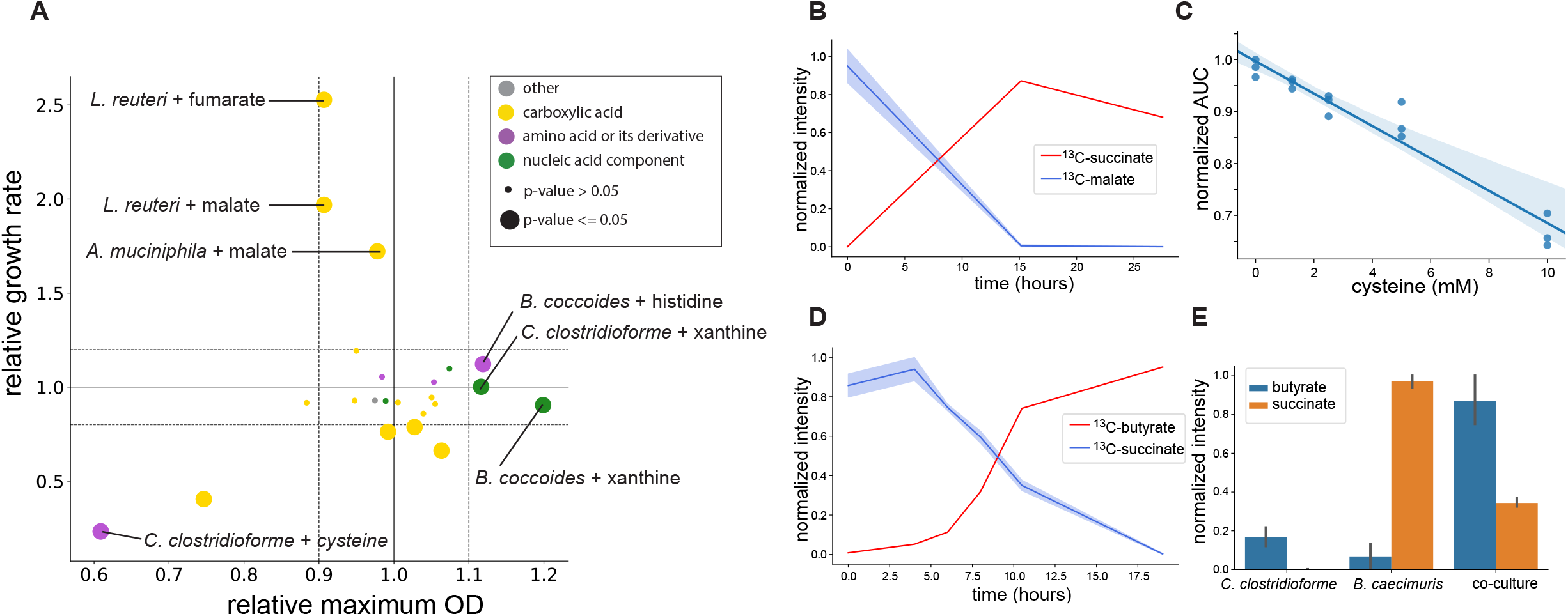
Supplementation and ^13^C-tracing experiments reveal impact of cross-feeding interactions. **A.** Impact of cross-fed nutrient supplementation on maximum OD and growth rate in mBHI supplemented with one metabolite of the indicated compound class (n=3 replicates per experiment). Values are shown relative to the growth rate and maximum OD obtained without supplementation. Color indicates metabolite class and dot size is proportional to the significance of the p-value (Student t-test). Dotted horizontal lines at 1.1 and 0.9 are shown for reference. **B.**Extracellular time course of fully ^13^C-labeled succinate and malate in *L. reuteri* mBHI cultures supplemented with ^13^C-malate. Shaded areas represent the standard deviation from the experiments (n=3 replicates) **C.**Normalized area under the OD curve of *C. clostridioforme* grown in mBHI supplemented with different concentrations of cysteine. Shaded areas represent the standard deviation from the mean (n= 3 replicates). **D.**Levels of ^13^C-succinate and ^13^C-butyrate over time when *C. clostridioforme* was grown in mBHI supplemented with ^13^C-succinate. Shaded areas represent the standard deviation from the mean (n = 3 replicates) **E.**Levels of succinate and butyrate in mono-cultures and co-culture of *C. clostridioforme* and *B. caecimuris* in GMM normalized to the maximum value across all experiments (n= 3 replicates).

The higher biomass yield is most likely explained by providing energetically expensive N-containing building blocks for biomass; *i.e*. synthesis of nucleotides and histidine requires several ATP per molecule. The dicarboxylic acids malate and fumarate could potentially function as anaerobic electron acceptors that are reduced to succinate, allowing bacteria to generate more ATP^36^. To confirm their metabolic fate, we supplemented *L. reuteri* and *A. muciniphila* mBHI cultures with fully ^13^C-labelled malate. Consistent with the hypothesis, both cultures consumed ^13^C-malate and secreted fully ^13^C-labeled succinate (Fig. 4B, Fig. S5A).

To assess the strong negative impact of cysteine, we supplemented *C. clostridioforme* cultures with fully ^13^C-labeled cysteine and identified pyruvate as the degradation product (Fig. S5B). The inevitable byproduct of this reaction is hydrogen sulfide, an important role in maintaining physiological homeostasis in the gut^42^, but toxic for host and bacteria at higher concentrations^43^. Indeed, increasing cysteine supplementation augmented the growth inhibition (Fig. 4C), suggesting hydrogen sulfide as the inhibiting agent. This inhibition might also be relevant *in vivo* because hydrogen sulfide is one of the four main gut microbiome gases^42^.

### Succinate cross-feeding is a main source of butyrate production in the OMM consortium

Since succinate accounted for the by far greatest mass flow of carbon in the OMM cross-feeding network (Fig. 3A) and is a known microbiota-derived metabolite with important roles in gut homeostasis, pathogen susceptibility, and inflammatory-related diseases^44^, we next investigated its metabolic fate. Under anaerobic conditions, succinate is generally considered as a reduced end product^36^ or a key intermediate of propionate synthesis by primary fermenters such as Bacteroides when CO_2_ is limiting^44,45^. To elucidate the fate of succinate in one of the two main consumers *C. clostridioforme*, we grew cultures in mBHI supplemented with fully ^13^C-labelled succinate and analyzed its intracellular metabolome during mid exponential growth at isotopic steady state by untargeted LC-MS. Among the fully labelled intracellular metabolites we found several intermediates of butyrate production (Fig. S4A, Table S12). Conversion of succinate to butyrate has been described for *C. kluyveri*^46^ and more recently also for the gut pathogen *C. difficile*^47^. In the latter case it was shown to constitute an important metabolic niche in the absence of other succinate consumers after antibiotic treatment^47^. While this conversion to butyrate does not produce ATP, it acts as an electron sink regenerating NAD+ from NADH (Fig. S4B). Consistently, we observed consumption of ^13^C-succinate and secretion of fully labelled ^13^C-butyrate (Fig. 4D), providing strong evidence for the operation of this pathway in *C. clostridioforme*.

To verify whether succinate cross-feeding actually occurs in co-culture, *C. clostridioforme* and *B. caecimuris* were grown in mono- and co-cultures. As described above (Fig. 1C; Fig. 3A), monocultures of *B. caecimuris* produced succinate, but no butyrate, while *C. clostridioforme* produced a small amount of butyrate (Fig. 4E). In co-culture, however, butyrate accumulated to much higher levels at the expense of succinate, demonstrating succinate cross-feeding and butyrate fermentation in *C. clostridioforme*. Overall these results show that the mice commensal *C. clostridioforme* is a butyrate producer, and succinate cross-feeding within the OMM consortium might be a relevant source of butyrate.

While this cross-feeding does not provide an *a priori* fitness benefit to *C. clostridioforme*,butyrate is a host-relevant metabolite that can be used as a carbon source by colonocytes^48^ and has anti-inflammatory properties^49^. Since succinate consumption improves gut colonization of the pathogen *C. difficile* in the presence of the dietary sugar sorbitol that requires NAD+ for its catabolism^47^, we grew *C. clostridioforme* in the rich gut microbiota medium with sorbitol or succinate and sorbitol as carbon sources. Akin to *C. difficile*, succinate availability improved the fitness of *C. clostridioforme*, albeit only to a small extent (Fig. S4C). Thus, we show that the benefits of succinate cross-feeding for *C. clostridioforme* are context dependent, and suggest that it might be a relevant interaction *in vivo* in the presence of the abundant diet-derived carbon source sorbitol.

## Discussion

Our results are based on an experimental approach that combines systematic *in vitro* cultivation in rich and spent media with dynamic exo-metabolomics to characterize metabolic fingerprints of species and infer potential metabolic interactions in microbial communities. Beyond the identification of producers of well-known gut microbiome fermentation products such as acetate, propionate, butyrate and lactate^40^, our systematic approach mapped 152 interactions in the recently introduced synthetic mouse gut consortium OMM^15^. As the major constituents with up to 50% of the consortium in the mouse colon^21^, the Bacteroidetes phylum representatives *B. caecimuris* and *M. intestinale* were the main providers, both in terms of numbers of compounds and in mass flow. The former dominated the C interaction network primarily with secretion of vast amounts of succinate, but produced also the electron acceptors fumarate and malate that could be used by several other species. While *M. intestinale* dominated the N interaction network with the secretion of large amounts of arginine, several other species contributed further N-containing compounds such as histidine and the nucleobases hypoxanthine and xanthine. As the member with the largest genome, *C. clostridioforme* was the by far most promiscuous consumer of metabolites in the consortium. *F. plautii* assumes a special role within the community in being one of the two main consumers of C in the form of succinate and a major producer of N-containing compounds.

While cross-feeding of carboxylic and amino acids was known to occur in the mammalian gut^50^, cross-feeding of nucleobases is to our knowledge a new observation. The extent of xanthine and hypoxanthine interactions in our consortium and nucleobase secretion by other gut microbes such as *E. coli*^51^ suggests that this cross-feeding might not be limited to the OMM consortium. Such purine metabolites were recently shown to affect host traits, including aging^52^, irritable bowel syndrome^53^ or maintenance of the mucus barrier function^54^. Supplementation experiments with the most abundant cross-fed metabolites demonstrated physiological benefits in several cases, even in rich complex medium. Generally, N cross-feeding improved biomass formation of *B. coccoides* and *C. clostridioforme*, in particular through histidine and xanthine. Although cross-fed at large quantities, arginine did not provide any fitness benefit to the consuming species, but led to the production of ornithine, a metabolite implied for mucosal health^39^ that can also induce the biosynthesis of enterobactins by *E. coli* during infection^55^.

In contrast to the N compounds, cross-feeding with C-containing compounds affected only the rate of biomass formation, presumably by allowing to produced more energy per carbon source. In particular, the anaerobic electron acceptors malate and fumarate greatly increased the specific growth rate of *L. reuteri* and *A. muciniphila*. In particular, the higher growth rate of *A. muciniphila* in the presence of malate might be a relevant interaction *in vivo*, since both *A. muciniphila* and *B. caecimuris* are the two most abundant members of the OMM consortium in the cecum and colon of adult mice^21^. Alternative electron acceptors like malate and fumarate have been shown to improve *in vivo* fitness of *E.coli*^35^ and the pathogen *Salmonella*^56^. Their production by *Bacteroides* species has been reported previously^57^ and is generally linked to the presence of Bacteroidetes^58^.

While physiological benefits to the consumer are a strong argument for relevant cross-feeding, a cross-feeding interaction might also be beneficial to the host. An example is succinate, that was reduced by *C. clostridioforme* to butyrate, a microbiome-derived metabolite shown to impact the host physiology as a carbon source, have anti-inflammatory function or act as a signaling compound^47^. Butyrate production from carbohydrates or organic acids has been described extensively^59^, in particular conversion of succinate to butyrate by *C. kluyveri*^46^ and more recently also for the gut pathogen *C. difficile*^47^. Through isotopic tracing and co-culture experiments with the major OMM succinate producer *B. caecimuris*, we demonstrated that also *C. clostridioforme* produces butyrate from succinate, which constituted the quantitatively largest cross-feeding flux in our consortium. This cross-feeding might be relevant *in vivo* not only as a host source of butyrate but also for depletion of the inflammatory succinate^60^ produced by other species. While our findings are limited to the OMM consortium, the prevalence of succinate producers such as Bacteroidetes in the gut microbiome indicates that succinate cross-feeding to butyrate might be a relevant source of butyrate in the gut microbiome.

The strongest negative fitness effect was seen for cysteine consumption by *C. clostridioforme*.While cysteine inhibition of amino acid biosynthesis has been reported for *E. coli*^61^, we confirmed its degradation to pyruvate through isotopic tracing. This degradation releases hydrogen sulfide, one of the four relevant microbiome-derived gases^42^. The origin of microbiome-derived hydrogen sulfide is often associated with the presence of sulfate-reducing bacteria, but members of this bacterial family are rather infrequent in the human microbiome^62^ and absent in the OMM consortium. Given that *C. clostridioforme* is an abundant OMM member *in vivo*^21^, it might play a key role in the degradation of cysteine and production of hydrogen sulfide in mice harboring the OMM consortium. Consistently, a recent study reported cysteine consumption in two other *C. clostridioforme* strains^58^, suggesting this species as a relevant cysteine consumer in the gut. Our findings are consistent with the recent notion that beyond dissimilatory hydrogen sulfide formation by sulfate reducers also cysteine catabolism is ubiquitous and an underestimated source of hydrogen sulfide human gut^63^.

Comprehensive characterization of the cross-feeding network suggests that the OMM consortium is highly connected at the metabolic level with distinct N and C interaction networks. Quantification and metabolic characterization of the main interactions revealed microbe-microbe interactions but also potential interactions with the host through metabolic end-products, including butyrate, hydrogen sulfide, and ornithine. While the here used *in vitro* experiments with complex mBHI and spent medium demonstrate only potential metabolic interactions in the gut, the recovery of known interactions, consistency with expected *in vivo* species abundance, and the demonstration of physiological relevance suggest that many of the cross-feeding interactions may also be relevant *in vivo*. Our findings show that many crucial metabolic features of the gut microbiota are represented within the OMM consortium and strengthen its relevance as a model for the mouse gut microbiota.

## Acknowledgement

We are grateful to Andrew Macpherson, Stephanie Ganal-Vonarburg and Jakob Zimmerman for providing the OMM species, continuous enlightening discussions, and the former for constructive feedback on the manuscript. This work was funded by the Swiss National Science Foundation (SNSF Sinergia CRSII5_177164) to US.

## Materials and Methods

### Chemicals and strains

All chemicals were purchased from Sigma-Aldrich. The OMM species were kindly provided by Andrew Macpherson^64^.

### Media preparation

Modified BHI (mBHI): 37 g l^-1^ Brain-Heart Infusion base, 5 mg l^-1^ hemin, 250 mg l^-1^ cysteine-HCl, 250 mg l^-1^ Na_2_S x 9 H_2_O, 0.5 mg l^-1^ menadione, and 0.25 g l^-1^ mucin from porcine stomach type II. Gut microbiota medium was prepared as previously^65^ except with no addition of SCFA and using sorbitol (0.5% w/v) as the only carbon source.

### Fresh and spent medium experiments

All strains were grown under anoxic conditions in an anaerobic chamber (Coy Laboratory Products Inc, MI, USA) filled with an anaerobic gas mix (5% (v/v) carbon dioxide, 5% (v/v) hydrogen, 90% (v/v) nitrogen) at 37°C. Overnight liquid cultures of 10 mBHI ml were prepared for each species from frozen stocks. For experiments with fresh mBHI, 50 ml mBHI were dispensed in 150 ml serum bottles (VWR International & Omnilab AG), sealed with a butyl rubber septum and inoculated from an overnight pre-culture to an initial OD of 0.05. For every consortium member, three to four replicates were incubated at 37°C and stirred at 300 rpm with a small cross-shaped stir bar (2 cm diameter). Aliquots for culture density measurements and metabolomics were withdrawn with a 1 ml syringe and 23G BD Precisionglide^®^ syringe needle through the rubber septum over the entire growth curve to capture lag, exponential, and stationary phase (Table S13).

Spent media were prepared from cultures in stationary phase. For this purpose, culture broth was dispended in 50 ml Falcon tubes and centrifuged at 3500 rpm for 10 minutes at 4°C, and the supernatant was filter sterilized (PES membranes with 0.22 μm pore size). Aliquots of spent media were stored at −20°C. Individual aliquots were thawed and equilibrated in the anaerobic chamber overnight to remove dissolved oxygen before utilization. For spent media experiments, Hungate tubes were filled with 10 ml of a 1:1 mixture of mBHI and spent medium from the specified species. For every spent medium experiment, Hungate tubes were inoculated in duplicate with 100 μl from an overnight mBHI pre-culture in stationary state to an approximate initial OD of 0.05. Optical density was followed over the course of the experiment by measuring it directly from the Hungate tube with a Biowave cell density meter CO8000 (VWR International). Similarly, to the fresh medium experiments, samples were collected over the course of the growth curve trying to capture the different phases of bacterial growth (Table S13).

### Mass spectrometry analysis

Aliquots for metabolomics analysis were prepared by centrifugation to separate cells from culture supernatant and stored at −80°C until further use as previously described^66^. For untargeted analysis, 40 x diluted and centrifuged samples were injected into an Agilent 6520 time-of flight mass spectrometer operated in negative mode at 2 Ghz EDR and with a mass / charge (m/z) range of 50-1000. The mobile phase was 60:40 (v/v) isopropanol:water and 1 mM NH4F at pH 9.0 for negative mode. For online mass axis correction mobile phases were supplemented with hexakis(1H, 1H, 3H-tetrafluoropropoxy)phosphazine and 3-amino-1-propanesulfonic acid for online mass correction. The injection sequence was randomized. Data was acquired in profile mode, centroided and analyzed with Matlab (The Mathworks, Natick). Missing values were filled by recursion in the raw data. Upon identification of consensus centroids across all samples, ions were putatively annotated by accurate mass and isotopic patterns. Starting from the a comprehensive list of bacterial metabolites database compiled by extracting the metabolites present in the KEGG genomes of gut bacteria ^67^. All formulas matching the measured mass within a mass tolerance of 0.003 Da were enumerated. As this method does not employ chromatographic separation or in-depth MS2 characterization, it is not possible to distinguish between compounds with identical molecular formula. The confidence of annotation reflects Level 4 but – in practice - in the case of intermediates of primary metabolism it is higher because they are the most abundant metabolites in cells.

Short chain fatty acids were quantified with via the 3-NPH derivatization method developed by Han and colleagues^68^. Briefly, 40 μl of 10-fold diluted samples were mixed with 20 μl 120 mM EDC-HCL-6% (v/v) pyridine solution and 20 μl of 200 mM 3-NPH HCL solution. Samples were incubated at 37°C for 30 minutes and diluted 25 times with 10% aqueous acetonitrile. Finally, samples were centrifuged 2 minutes at 3500 rpm and the clear supernatant was sued for analysis. Samples were measured with the same MS system as described above. Chromatographic separation was performed on a 50 x 2.1mm, 130 Å, 1.7 μm Acquity UPLC BEH C18 column (Waters) using a mobile phase A: H_2_O, 0.1% formic acid and B: acetonitrile, 0.1% formic acid. An injection volume of 2 μl was used and elution was achieved using the following gradient: initial conditions, 83% A, 1100 μl/min; 0.2 min 83% A; 1.9 min 82% A; 2.8 min 60% A; 3.0 min 0% A; 3.50 min 0% A; 3.51 min 83% A. Online mass calibration was performed using a second spray needle and a constant flow (5 ul/min) of reference solution containing purine and hexakis (1H, 1H, 3H -tetrafluoropropoxy)phosphazine (HP-0921, Agilent Technologies). Compounds were identified based on the retention time of chemical standards and their accurate mass (tolerance 20 ppm). The MassHunter Quantitative Analysis Software (Agilent, version 7.0) was used for peak integration.

Quantitative measurement of selected metabolites was performed with liquid chromatography coupled to MS. Chromatographic separation via hydrophilic interaction (HILIC) was performed on an AdvanceBio MS Spent Media column (50 x2.1mm, Agilent technologies) using a mobile phase A: H_2_O, 10 mM ammonium acetate, pH 9.0 and B: Acetonitrile, 10 mM ammonium acetate, pH 9.0. Samples were prepared as above and diluted 5 times in 50:50 water: acetonitrile. One μl of 100-fold diluted sample was injected and elution was achieved using the following gradient: initial conditions, 5% A, 1000 μl/min; 0.25 min 5% A; 0.75 min 50% A; 1.0 min 65% A; 1.25 min 65%; 1.26 min 95%; 2.25 min 95% A. The qTOF (Agilent 6520) was operated in negative mode at 2 GHz extended dynamic range and with mass/ charge (m/z) range of 50-1000 and the following source parameters: VCap: 3500 C, nozzle voltage: 2000 V, gas temp: 325°C m drying gas 5 l/min, nebulizer 30 psig. Online mass calibration was performed using a second spray needle and a constant flow (5 ul/min) of reference solution containing purine and hexakis (1H, 1H, 3H -tetrafluoropropoxy)phosphazine (HP-0921, Agilent Technologies). Compounds were identified based on the retention time of chemical standards and their accurate mass (tolerance 20 ppm). The MassHunter Quantitative Analysis Software (Agilent, version 7.0) was used for peak integration and quantification was performed in the software based on a calibration curve of chemical standards.

Cysteine and cystine quantification was performed with liquid chromatography coupled to MS using an 5500 QTRAP triple quadrupole mass spectrometer in positive mode and MRM scan type (AB SCIEX, Foster City, CA). Separation was performed using a HILIC Plus RRHD column (1.8 μm, 2.1×100 mm, Agilent technologies) using a mobile phase A: H_2_O with 0.1 % formic acid (v/v) 10 mM ammonium formate and B: Acetonitrile with 0.1% formic acid (v/v). Five μl of 80x diluted sample were injected and elution was achieved using the following gradient initial conditions, 10% A, 400 μl/min; 2.0 min 60% A; 3.0 min 60% A; 5.0 min 10% A; 6.0 min 10% A. Data acquisition was performed with Analyst 1.7.1 software (Sciex, Darmstadt, Germany) and peak integration was performed using in-house software. To account for the oxidation over time of cysteine in spent media^69^, cysteine was quantified by adding the concentration of cysteine plus two times the concentration of cystine.

For labelling experiments, 100-fold diluted and centrifuged samples were injected into an Agilent 6546 time-of flight mass spectrometer operated in negative mode with a mass / charge (m/z) range of 50-1000. Chromatographic separation was performed on a 30 x 2.1mm, 1.7 μm Acquity UPLC BEH C18 column (Waters) using a mobile phase A: H2O, 0.1% acetic acid and B: Methanol, 0.1% acetic acid. An injection volume of 2 μl was used and elution was achieved using the following gradient: initial conditions, 83% A,1100 μl/min; 0.2 min 83% A; 1.9 min 82% A; 2.8 min 60%A; 3.0 min 0% A; 3.50 min 0% A; 3.51 min 83% A. Online mass calibration was performed using a second spray needle and a constant flow of reference solution containing purine and hexakis (1H, 1H, 3H -tetrafluoropropoxy)phosphazine (HP-0921, Agilent Technologies). After processing of raw data as previously described^24^, m/z features (ions) were annotated by matching them to the accurate mass-to-sum formulas of a comprehensive list of bacterial metabolites database with 0.001 Da mass accuracy assuming single deprotonation [M-H. Notably, this metabolomics method cannot distinguish between isobaric compounds, e.g. metabolites having identical m/z values (e.g. leucine vs. isoleucine).

### Identification of consumed and secreted metabolites

Increasing and decreasing metabolites were identified from the untargeted metabolomics dataset as annotated ions with significant correlation to the OD of the bacteria over time (Pearson correlation coefficient >0.7, p-value<0.05) or significant goodness of linear or exponential fit (R2>0.7, p-value <0.05). The latter accounts for metabolites that might be exhausted before the end of the growth experiment or constantly produced throughout the experiment. Furthermore, the maximum fold change between the initial time point and any other point had to be higher than −1.37 for consumed metabolites and higher than 1.20 for a secreted metabolite. These thresholds were determined based on an average fold change observed for every annotated ion in mBHI in a dilution series. In brief, 20-to 720-fold dilutions of mBHI medium were measured with FIA-MS in triplicate. Since FIA-MS is sensitive to matrix effects, ion count changes cannot be directly translated into an equivalent metabolite change.

To remove background ions that are not derived from mBHI the above thresholds were determined focusing only with annotated metabolites that had a high significant negative correlation to the dilution factor (Pearson correlation < −0.75, Fig. S6A, B). To determine average fold changes of consumed metabolites, we used measurements of the annotated metabolites in 40-80-fold dilutions. To determine average fold changes of secreted metabolites, we used measurements of annotated metabolites in 20-to 40-fold dilutions. Average fold changes were 1.37 and 1.20 for consumed and secreted metabolites, respectively (Fig. S6C).

### *In vitro* supplementation experiments

Overnight liquid cultures of 5 ml of mBHI were prepared for each species from frozen stocks. Solutions of supplemented metabolites were prepared in deionized water, titrated to pH 7, filter sterilized (0.22 μm) and, stored at −20°C. Stock solutions of supplements were thawed and equilibrated overnight in the anaerobic chamber the day before the start of the experiment. Species were then grown anaerobically in in triplicates in 10 ml Hungate tubes consisting of 90% (v/v) mBHI and 10% (v/v) supplement at 10 mM. Growth curves were acquired by following OD with a Biowave cell density meter CO8000 (VWR International). Growth rates were inferred by fitting growth-curves to OD as a function of time with a four-parameter logistic function was used^70^.

### Isotopic tracer experiments

Overnight liquid cultures of 5 ml of mBHI were prepared for each species from frozen stocks. Solutions of labelled supplemented metabolites were prepared in deionized water, titrated to pH 7, filter sterilized (0.22 μm) and, stored at −20°C. Stock solutions of supplements were thawed and equilibrated overnight in the anaerobic chamber the day before the start of the experiment. Species were then grown anaerobically in triplicates in 10 ml Hungate tubes consisting of 90% (v/v) mBHI and 10% (v/v) supplement at 10 mM except for ^13^C,^15^N-L-Cysteine that was supplemented at 1 mM. Growth curves were acquired by following OD with a Biowave cell density meter CO8000 (VWR International). Supernatant samples were collected and processed as explained above. Intracellular samples were obtained at mid exponential phase (OD of 0.4-0.6) and extracted as previously described^71^. In brief, 2 ml aliquots of cell culture were filtered by vacuum filtration on a 0.45-μm filter. On the filter, cells were washed with 2 ml of pre-warmed ammonium carbonate buffer at pH 7.2. Filters with cultured cells were immediately transferred for extraction into 2:2:1 acetonitrile:methanol:water solution at −20°C. Cells were extracted for at least 2 hours and centrifuged at 14000 g at 4°C for 20 minutes to remove cell debris. Supernatants were dried at 0.12 mbar and resuspended in 100 μl of deionized water.

### OMM members metabolic potential assessment

For each of the ten cultivated strains of the OMM consortium except *L. reuteri*, *E. faecalis* and *B. longum*, a list of KEGG orthologues (KO) for the protein-coding genes were obtained from the KEGG genome database ^72^. For *L. reuteri, E*. faecalis and *B. longum* the KEGG Automatic Annotation Server(KAAS) with the Blast default settings^73^ was used to obtain a list of KEGG orthologues. Gut metabolic module (GMM) detection was performed as previously described^67^. In brief, GMM presence/absence was identified with a detection threshold of more than 50% coverage.

## Data availability

Raw mass spectrometry data and growth physiology data can be downloaded from https://www.ebi.ac.uk/biostudies/, accession code S-BSST686.

---

**Supplementary Figure 1 A.** Metabolites consumed (blue) by more than half of the OMM members. Metabolites that were not detected as abundant in fresh mBHI are shown in bold letters. Intensities are scaled to +/− 1 by dividing each metabolite by the maximum observed change in abundance in all species. **B.** Metabolites that were uniquely secreted by one member of the consortium. Metabolites that were not detected as abundant in fresh mBHI are shown in bold letters. Alternative annotation within the mass tolerance of 3 mDa are indicated where present. **C.** Upset plot of consumed metabolites by the OMM species. **D.** Upset plot of produced metabolites by the OMM species.

**Supplementary Figure 2** Consumption (blue) and secretion (red) of relevant metabolite classes by OMM species. Consumption or secretion was inferred by comparing late stationary phase concentrations in spent media to fresh mBHI medium concentrations, scaled to +/− 1 by dividing the relative concentration of each metabolite by the maximum observed change in abundance in all species. **A.** Nucleic acid profiles determined by untargeted metabolomics analysis with FIA-MS. **B.** Short chain fatty acid profiles determined by 3-NPH derivatization^68^ coupled with targeted metabolomics. **C.** Vitamin and cofactor profile determined by untargeted metabolomics analysis with FIA-MS. **D.** Amino acid profile determined by HILIC LC-MS. Asterisk’s indicate species with the ability to degrade an amino acid via Stickland fermentation that consumed the respective amino acid.

**Supplementary Figure 3.** Relative abundance of OMM members in mice colonized with the OMM consortium. Relative abundance in three different locations was determined by metagenomics sequencing. Data was taken from Yilmaz *et al*^21^.

**Supplementary Figure 4. A.** Succinate to butyrate conversion pathway proposed in *C. clostridioforme*.Metabolites highlighted in bold were detected as fully labelled intracellularly when *C. clostridioforme* was supplemented with ^13^C-succinate **B.** Growth of *C. clostridioforme* in gut microbiota medium with either sorbitol or sorbitol and succinate. Shaded areas represent the standard deviation from the mean (n=3).

**Supplementary Figure 5. A.** Time course of extracellular fully ^13^C-labeled succinate and malate in *A. muciniphila* in mBHI cultures supplemented with ^13^C-malate. Shaded areas represent the standard deviation from the experiments (n=3). **B.** Intracellular fully labelled fraction of pyruvate in *C. clostridioforme* when grown with ^13^C-cysteine or with ddH_2_O (n=3).

**Supplementary Figure 6. A.** Distribution of Pearson correlation coefficients between dilution factor and ion intensity of the metabolites annotated in mBHI via FIA-MS. **B.** Normalized ion intensity over the dilution factor for metabolites with a Pearson correlation coefficient below −0.75. Shaded areas represent the standard deviation from the mean (n=3) **C.** Average fold change for annotated ions that had a Pearson correlation coefficient below −0.75 from 20-to 40-fold dilution (increase) and from 40-to 80-fold dilution (decrease).

