## Supplementary figures for "Distinct N and C cross-feeding networks in a synthetic mouse gut consortium"

### Supplementary Figure 1

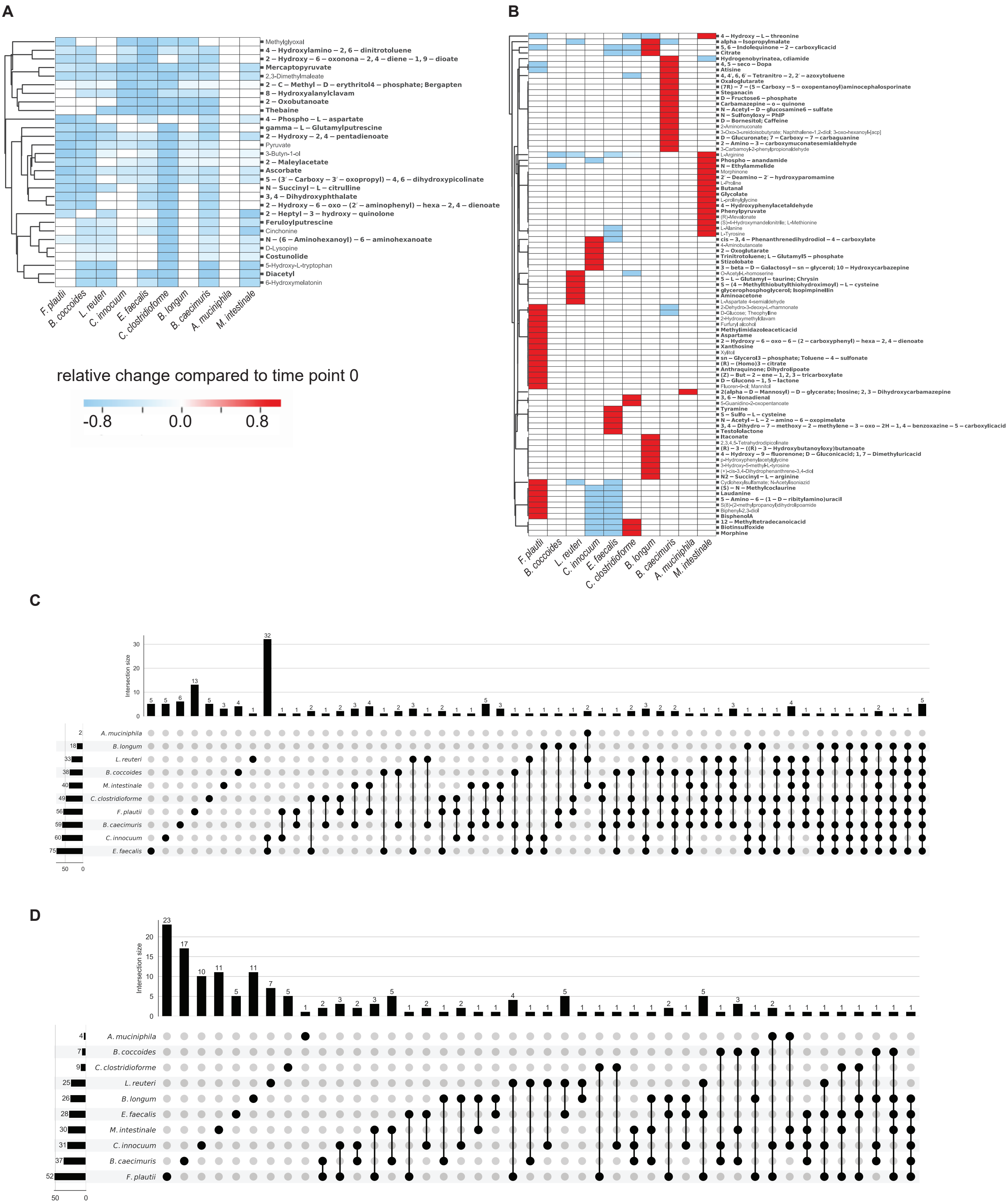

### Supplementary Figure 2

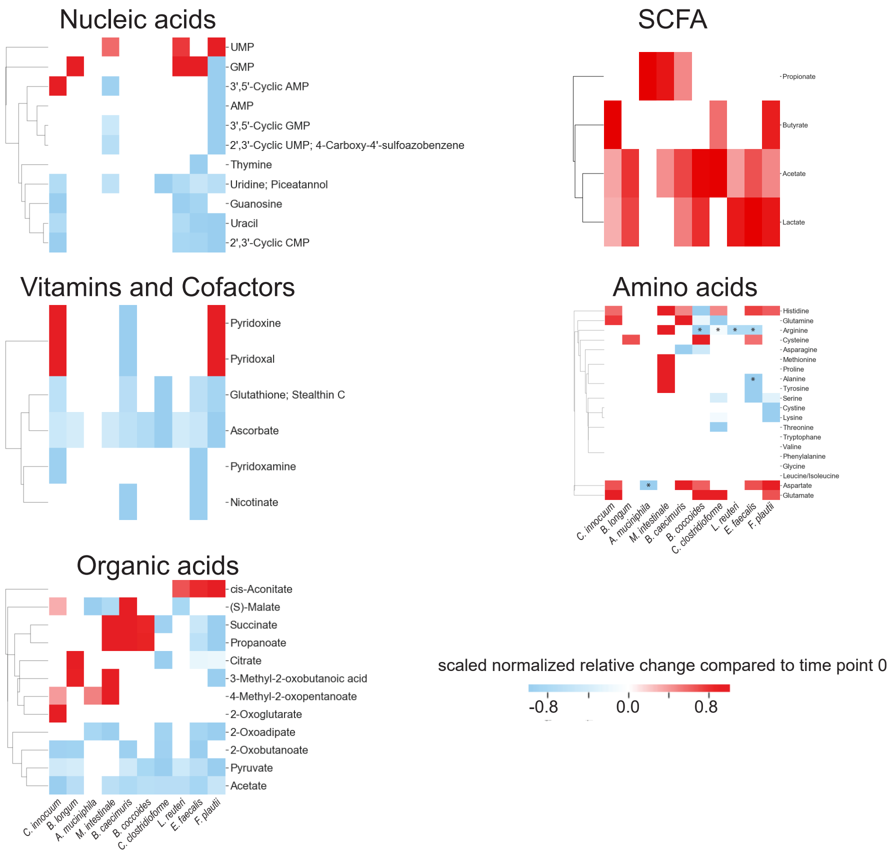

### Supplementary Figure 3

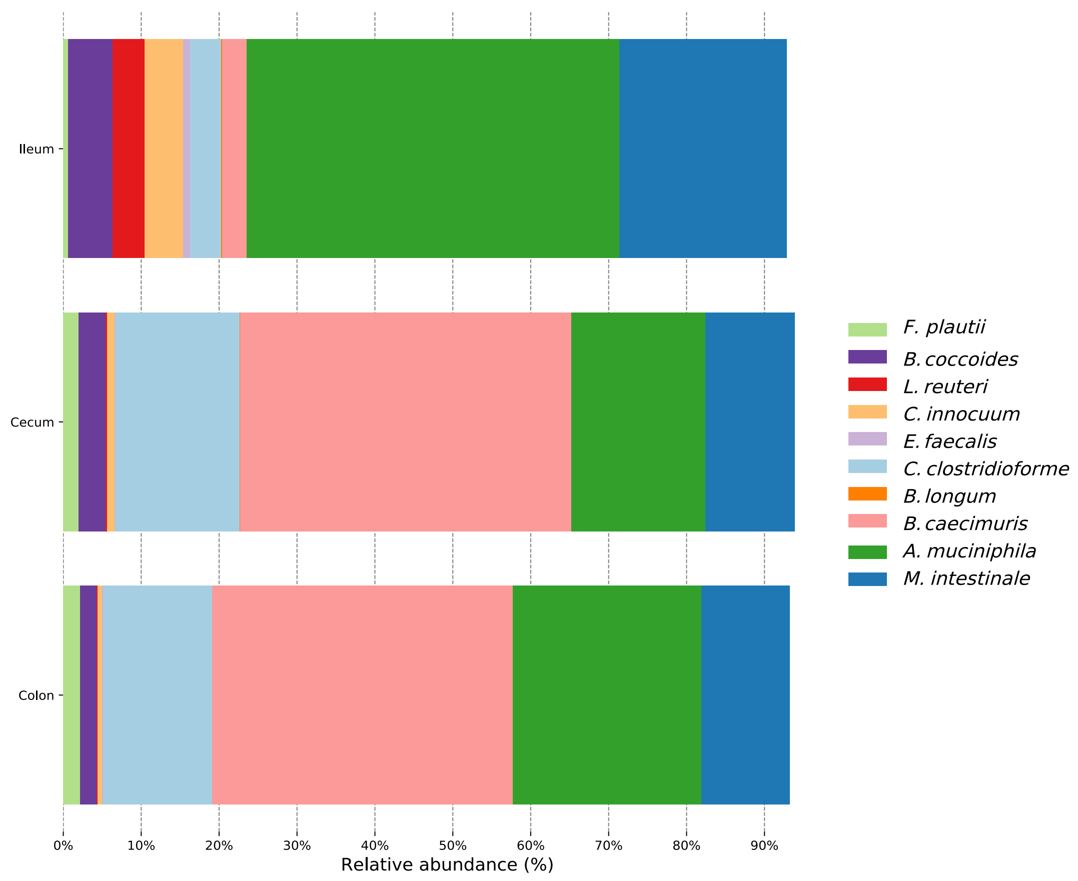

### Supplementary Figure 4

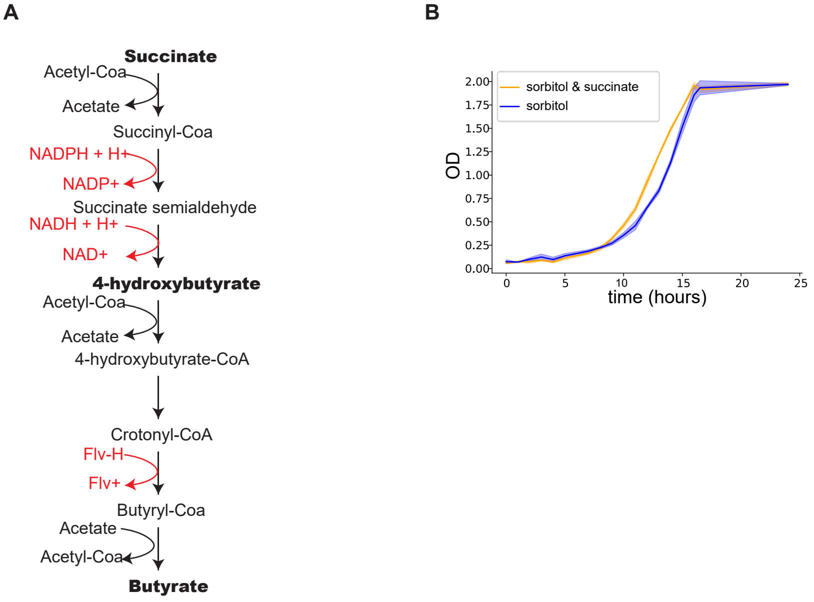

### Supplementary Figure 5

**A**

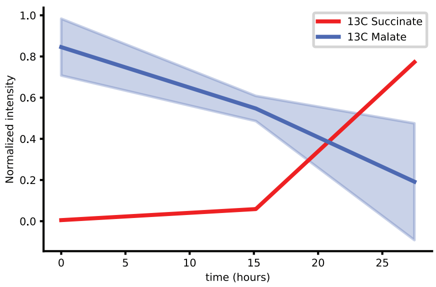


**B**

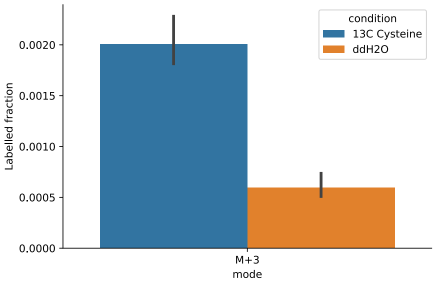

### Supplementary Figure 6

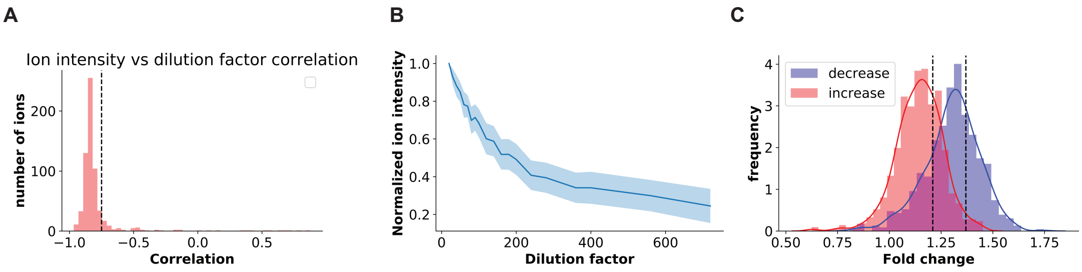
